# Disruption of Z-RNA–binding of ADAR1 induces Aicardi-Goutières syndrome–like encephalopathy in mice

**DOI:** 10.1101/2020.12.16.422984

**Authors:** Taisuke Nakahama, Yuki Kato, Toshiharu Shibuya, Jung In Kim, Tuangtong Vongpipatana, Hiroyuki Todo, Yanfang Xing, Yukio Kawahara

**Affiliations:** Department of RNA Biology and Neuroscience, Osaka University, Suita, Osaka 565-0871, Japan; Integrated Frontier Research for Medical Science Division, Institute for Open and Transdisciplinary Research Initiatives (OTRI), Osaka University, Suita, Osaka 565-0871, Japan; the Genome Editing Research and Development Center, Graduate School of Medicine, Osaka University, Suita, Osaka 565-0871, Japan

**Keywords:** ADAR1, AGS, ISG, MDA5, RNA editing, Z-RNA

## Abstract

ADAR1 p150 is an enzyme responsible for adenosine-to-inosine RNA editing. Deletion of ADAR1 p150 results in embryonic lethality with a type I interferon (IFN) signature, caused by aberrant MDA5 sensing unedited transcripts. ADAR1 p150 contains a unique Z-DNA/RNA–binding domain α (Zα); however, the role of this domain remains unknown. A mutation has been identified in this domain in patients with Aicardi-Goutières syndrome (AGS), an inherited interferonopathy, suggesting an essential role in avoiding MDA5 activation. Here, we show that a mutation in the Zα domain reduces the editing activity of ADAR1 p150 by comparing activity between wild-type and mutated isoforms expressed in *Adar1/Adar2* knockout cells. Furthermore, we created Zα domain–mutated knock-in mice, which displayed severe growth retardation with abnormal organ development, including AGS-like encephalopathy with a type I IFN signature. These abnormalities were ameliorated by the concurrent deletion of MDA5. Collectively, Z-RNA–recognition contributes to ADAR1 p150–mediated RNA editing, which prevents MDA5 activation.

## INTRODUCTION

Adenosine (A)-to-inosine (I) RNA editing is a widely conserved post-transcriptional RNA modification essential for survival in mammals (Bass, 2002; Heraud-Farlow and Walkley, 2020; Nishikura, 2010). Adenosine deaminases acting on RNA (ADARs) are enzymes responsible for this modification with two active ADARs, ADAR1 and ADAR2, identified so far in mammals (Nishikura, 2016; Slotkin and Nishikura, 2013; Zipeto et al., 2015). Given that ADARs contain double-stranded (ds)RNA-binding domains (dsRBDs) (**Figure 1A**), A-to-I RNA editing occurs in dsRNA structures, which are preferentially formed by inverted repetitive sequences, such as short interspersed elements (SINEs), located in introns and 3’ untranslated regions (UTRs) of mRNA (Bazak et al., 2014; Levanon et al., 2004; Ramaswami et al., 2012).

**Figure 1.**
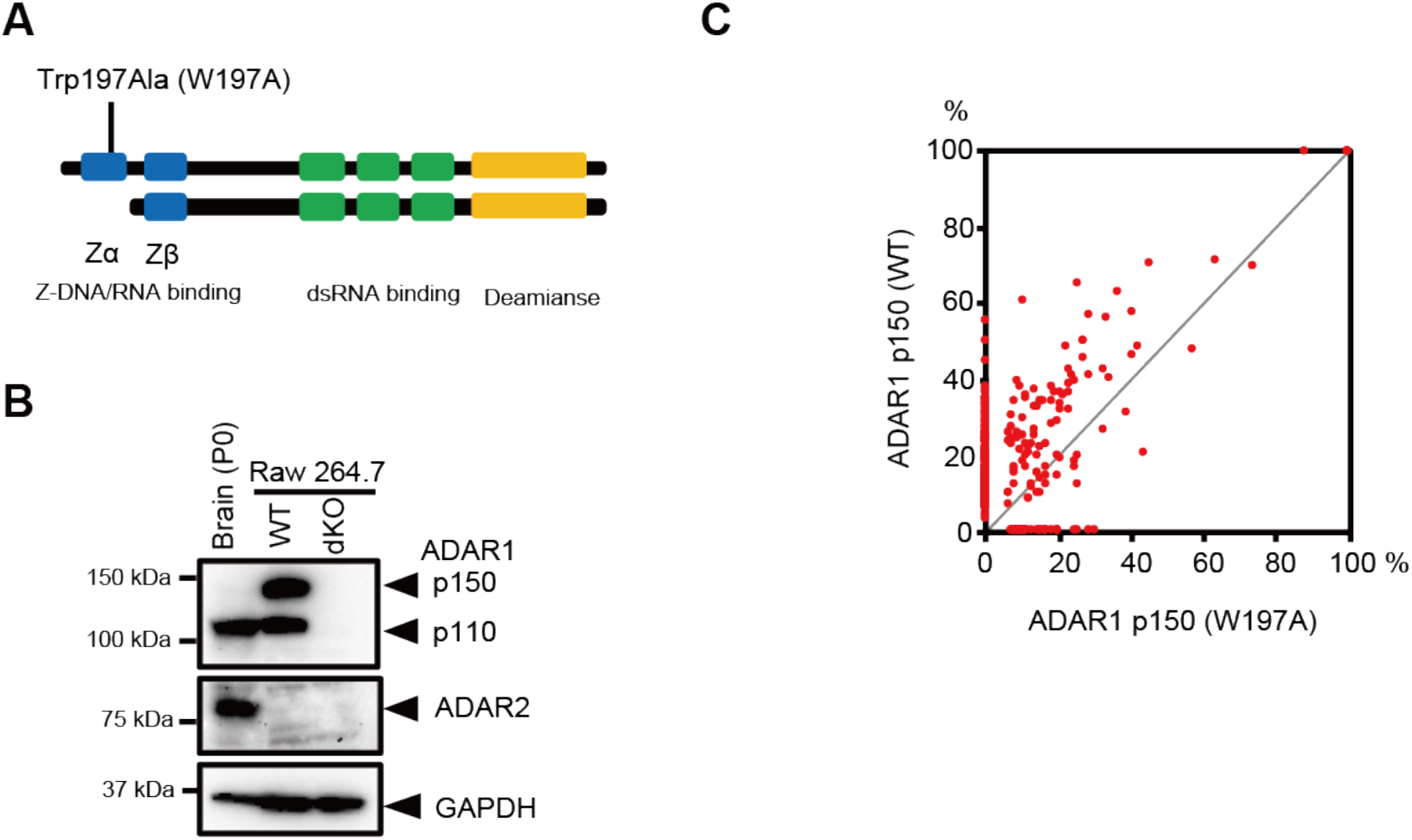
Comparison of the editing activity between wild-type and Zα domain– mutated ADAR1 p150 isoforms. **(A)** Schematic diagram of mouse ADAR1 p110 and p150 isoforms. One Z-DNA/RNA binding domain (Zβ; blue), three dsRNA binding domains (green) and a deaminase domain (yellow) are shared by both isoforms, while the N-terminal Zα domain is unique to the ADAR1 p150 isoform. Tryptophan-to-alanine substitution at the position of amino acid 197 (W197A) in the Zα domain leads to a loss of binding capacity to Z-DNA/RNA. **(B)** Immunoblot analysis of ADAR1 p110, ADAR1 p150, ADAR2, and GAPDH protein expression in wild-type (WT) and *Adar1/Adar2* double knockout (dKO) Raw 264.7 cell lines. The expression of these proteins in the brain at post-natal day 0 (P0) is shown as a reference. **(C)** Editing ratios of the sites identified are compared between WT and Zα domain–mutated (W197A) ADAR1 p150 isoforms expressed in *Adar1/Adar2* dKO Raw 264.7 cells. Values are displayed as the mean of values from two independent experiments. We considered only sites covered with more than five reads in all samples.

ADAR1 is composed of two isoforms: longer ADAR1 p150 and shorter p110, which are transcribed from the same gene locus using different promoters (George et al., 2005; George et al., 2008; George and Samuel, 1999; Kawakubo and Samuel, 2000). ADAR1 p110 is localized in the nucleus and highly expressed in the mouse brain, whereas interferon (IFN)-inducible ADAR1 p150 is predominantly localized in the cytoplasm and abundant in lymphoid organs such as the thymus and spleen (Costa Cruz et al., 2020; Huntley et al., 2016; Nakahama et al., 2018; Tan et al., 2017). Of note, although ADAR1 p110 and p150 isoforms share the same Z-DNA/RNA binding domain β (Zβ), three dsRBDs, and a deaminase domain, p150 isoform specifically contains a Z-DNA/RNA binding domain α (Zα) in the N-terminus (Patterson and Samuel, 1995) (**Figure 1A**). This domain was originally identified as a domain that can bind to lefthanded Z-DNA (Herbert et al., 1995; Herbert et al., 1997) and was later proven to bind to Z-RNA (Brown et al., 2000; Herbert, 2019; Placido et al., 2007). However, although this domain has been shown to modulate RNA editing activity *in vitro* (Koeris et al., 2005), the functional role of this domain in RNA editing *in vivo* and the endogenous sequences that allow Z-RNA binding to this domain remains unidentified.

*Adar1* knockout (KO; *Adar1^-/-^*) and *Adar1* knock-in (KI) mice harboring the editing-inactive E861A point mutation (*Adar1^E861A/E861A^* mice) show embryonic lethality due to massive apoptosis accompanied by an overproduction of type I IFN and the resultant upregulated expression of IFN-stimulated genes (ISGs) (Hartner et al., 2004; Hartner et al., 2009; Liddicoat et al., 2015; Wang et al., 2000; Wang et al., 2004). This lethality can be rescued by the concurrent deletion of either *Ifih1*-encoded melanoma differentiation–associated gene 5 (MDA5), a cytosolic sensor for viral dsRNA, or mitochondrial antiviral signaling protein (MAVS), an adaptor protein downstream of MDA5 (Liddicoat et al., 2015; Mannion et al., 2014; Pestal et al., 2015). This evidence suggests that ADAR1-mediated RNA editing alters dsRNA structure to escape MDA5-mediated recognition of endogenous dsRNA as non-self, leading to activation of the innate immune system. In particular, ADAR1 p150 most likely exerts this function, given that *Adar1 p150*-specific KO (*Adar1 p150^-/-^*) mice also display similar embryonic lethality, which is rescued by the concurrent deletion of MAVS (Pestal et al., 2015; Ward et al., 2011). Therefore, it is naturally expected that the isoform-specific Zα domain may contribute to RNA editing of certain specific targets, which is required in order to avoid MDA5 activation. Of note, *ADAR1* mutations can cause Aicardi–Goutières syndrome (AGS), which is characterized as a type I interferonopathy affecting multiple organs, especially brain and skin, with upregulated expression of ISGs (Crow et al., 2015; Rice et al., 2012). Although most mutations have been identified in the deaminase domain, most likely reducing RNA editing activity, a point mutation is also identified at the position of amino acid 193 in the Zα domain, which converts proline (P193) to alanine (A). Using d(CG)-repeat DNA sequences that can form Z-DNA thermodynamically *in vitro*, it was shown that W195 in the β sheet of the Zα domain is the most critical site for interaction with Z-DNA (Schade et al., 1999). In addition, N173 and Y177 in the a helix of this domain are essential, whereas P193 also plays a supportive role in this interaction (Lee et al., 2018). Therefore, the reduced binding capacity of the mutated Zα domain to Z-RNA might lead to reduced RNA editing at certain ADAR1 p150–specific sites, which plays a central role in the pathogenesis of AGS. However, the physiological significance of the Zα domain remains to be examined *in vivo*.

In this study, we compared the editing frequency between wild-type ADAR1 p150 (WT) and mutated ADAR1 p150 (W197A) by expressing such expression constructs in *Adar1/Adar2* double KO (dKO) Raw 264.7 cells. W197 in mouse ADAR1 corresponds to W195 in humans. This analysis revealed that the editing ratio at ~85% of the sites in ADAR1 p150 (W197A)-expressing cells was lower than that in ADAR1 p150 (WT)-expressing cells. In contrast, no editing was observed when we expressed ADAR1 p150 (mut_dsRBDs), in which the Zα domain was intact, whereas the three dsRBDs were mutated to abolish binding to dsRNA through these domains (Valente and Nishikura, 2007). These results suggest that the presence of the Zα domain alone is not sufficient to induce RNA editing but that this domain contributes to efficient RNA editing at certain specific sites. We further generated *Adar1^W197A/W197A^* mice to elucidate the importance of Zα domain–mediated Z-RNA recognition *in vivo. Adar1^W197A/W197A^* mice displayed severe growth retardation after birth with abnormal development of multiple organs, including the spleen and colon. Of note, we observed that malformation of the brain was accompanied by gliosis, which is reminiscent of encephalopathy found in patients with AGS. The expression of ISGs was upregulated in the affected organs, which was restored by the concurrent deletion of MDA5. The expression of mutated ADAR1 p150 was increased in a feedback manner in *Adar1^W197A/W197A^* mice, which led to increased RNA editing at a subset of sites that most likely did not require the Zα domain. However, this compensatory upregulation of ADAR1 p150 expression could not ameliorate an inflammatory ISG signature. These findings suggest that the Zα domain–mediated tuning of RNA editing is essential for escaping MDA5 sensing of certain endogenous dsRNAs as non-self.

## RESULTS AND DISCUSSION

### W197A mutation in the Zα domain of ADAR1 reduces RNA editing activity

To examine the contribution of the Zα domain to ADAR1 p150–mediated RNA editing, we initially created an *Adar1/Adar2* dKO cell line by using mouse macrophage-like Raw 264.7 cells in which both ADAR1 p110 and p150 were abundantly expressed (**Figure 1A, B**). In this cell line, ADAR2 was barely detectable; however, we deleted both *Adar1* and *Adar2* genes by genome editing using a CRISPR/Cas9 system to exclude the possibility that a subtle amount of ADAR2 contributed to the RNA editing of certain sites (**Figure 1B**). A plasmid containing enhanced green fluorescent protein (EGFP)-tagged wild-type ADAR1 p150 or its mutants was then transfected into dKO cells. Twenty-four hours after transfection, cells displaying GFP fluorescence within the same range of intensity were sorted to adjust the expression of each ADAR1 p150 protein. Subsequently, RNA editing sites were analyzed by RNA-sequencing (RNA-seq). Although RNA editing sites are predominantly located in introns, which are generally spliced out in the nucleus (Hsiao et al., 2018; Licht et al., 2019), the ADAR1 p150 isoform is mainly localized in the cytoplasm. Therefore, we chose poly(A) RNA-seq to sensitively capture the difference in RNA editing activity by reducing the number of intronic sites. We showed that none of the sites identified in wild-type Raw 264.7 cells were detectable in dKO cells except for one site (*Sfi1*) which was verified to be a false-positive by direct Sanger sequencing (**Supplementary Table S1**). We first examined whether the Zα domain–mediated binding to dsRNA induces RNA editing by expressing ADAR1 p150 (mut_dsRBDs) in which the Zα domain was intact whereas the three dsRBDs were mutated to abolish binding to dsRNA through these domains. However, we did not detect any editing sites (**Supplementary Table S1**), which suggests that the presence of the Zα domain alone is not sufficient to induce RNA editing. Next, we compared the editing activity of ADAR1 p150 (WT) and ADAR1 p150 (W197A) by expressing these in dKO cells. The W197A mutation has been reported to completely abolish binding to Z-DNA (Schade et al., 1999). This analysis demonstrated that the editing ratio at ~85% of the sites in ADAR1 p150 (W197A)–expressing cells was lower than that in ADAR1 p150 (WT)–expressing cells (**Figure 1C**). These results suggest that the recognition of Z-RNA by the Zα domain contributes to ADAR1 p150–mediated RNA editing at certain sites.

### *Adar1^W197A/W197A^* mice display MDA5-depedendent AGS-like encephalopathy

To determine the physiological role of Z-RNA recognition by ADAR1 p150 *in vivo*, we intended to insert W197A or N175A/W197A point mutation(s) into the mouse *Adar1* gene to establish a KI model by genome editing using a CRISPR/Cas9 system. For this purpose, mouse embryos obtained from *Ifih1^-/-^* mice were used to avoid possible embryonic lethality, as observed in *Adar1^E861A/E861A^* and *Adar1 p150^-/-^* mice (Liddicoat et al., 2015; Pestal et al., 2015; Ward et al., 2011). We successfully obtained both *Adar1^W197A/W197A^* and *Adar1^N175A·W197A/N175A·W197A^* mice in an *Ifih1^-/-^* background. Although *Adar1^W197A/W197A^ Ifih1^-/-^* mice were slightly but significantly smaller than *Adar1^+/+^* mice, no other abnormal phenotypes were observed (**Figure 2A-C**). Therefore, we next restituted the *Ifih1* gene by crossing with WT mice to obtain *Adar1^W197A/+^* mice. These heterozygous mutant mice had a normal appearance and development. Therefore, by crossing heterozygous mutant mice, we generated *Adar1^W197A/W197A^* mice. Surprisingly, although their appearance seemed normal at birth, *Adar1^W197A/W197A^* mice displayed severe growth retardation during post-natal development, with approximately half of these mice dying by up to 6 weeks of age (**Figure 2A–2C**). Furthermore, *Adar1^W197A/W197A^* mice were too small and lean to survive without maternal care even after a normal weaning period of 3 – 4 weeks. These abnormalities were also observed in *Adar1^N175A·W197A/N175A·W197A^* mice, and therefore we used *Adar1^W197A/W197A^* mice for subsequent experiments.

**Figure 2.**
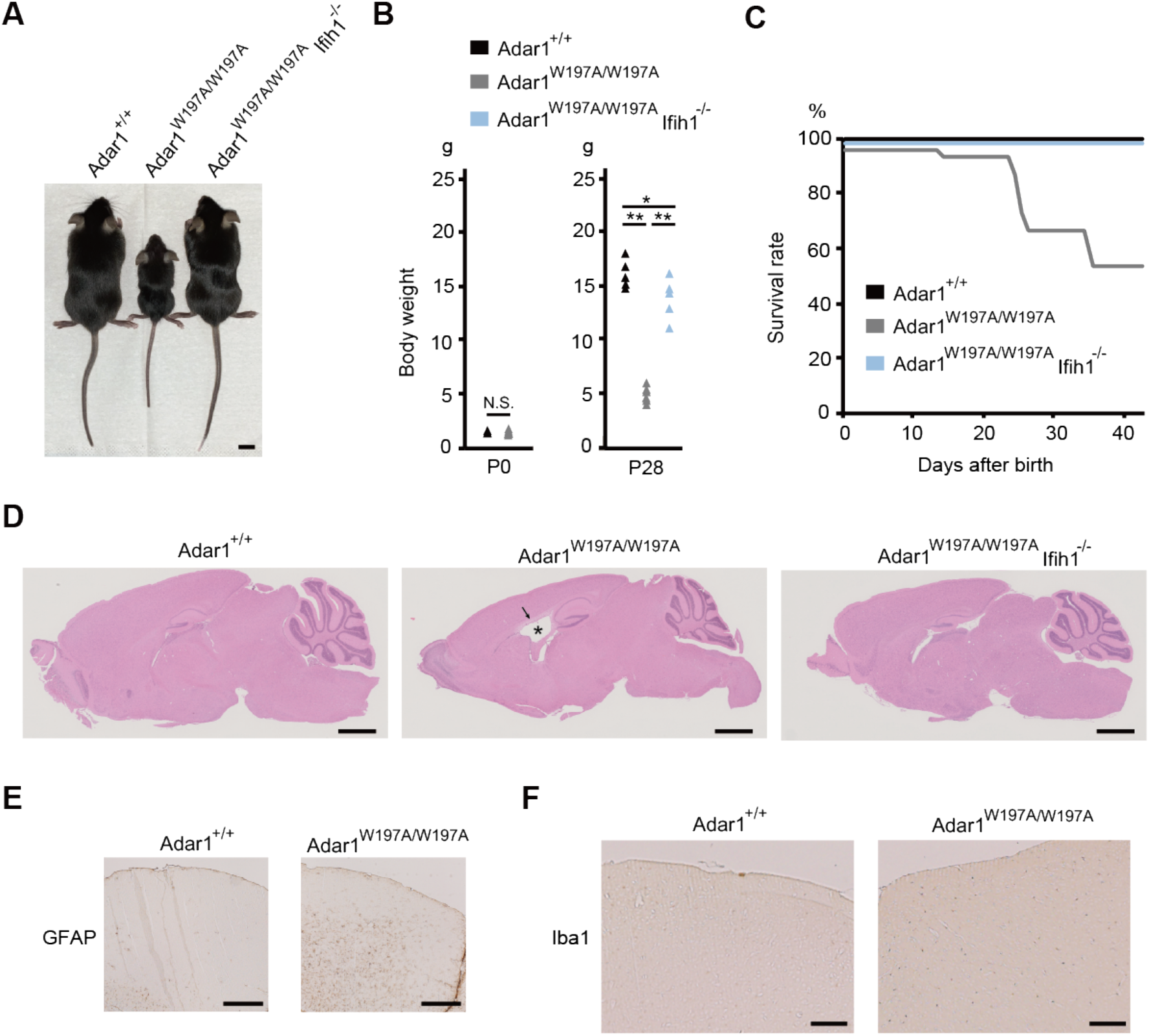
*Adar1^W197A/W197A^* mice display severe growth retardation. **(A)** Representative image of *Adar1^+/+^, Adar1^W197A/W197A^*, and *Adar1^W197A/W197A^ Ifih1^-/-^* mice at six weeks of age. Scale bar, 1 cm. **(B)** Body weights at P0 (left panel) and P28 (right panel) are compared between *Adar1^+/+^* (n = 3 at P0 and 5 at P28), *Adar1^W197A/W197A^* (n = 7 at P0 and 6 at P28), and *Adar1^W197A/W197A^ Ifih1^-/-^* (n = 5 at P28) mice. Mann–Whitney *U*-test, *\*p* < 0.05, \*\**p* < 0.01, N.S., not significant. **(C)** Survival curve of *Adar1^+/+^* (n = 9), *Adar1^W197A/W197A^* (n = 15), and *Adar1^W197A/W197A^ Ifih1^-/-^* (n = 8) mice. (**D**) Representative images of hematoxylin and eosin (HE) staining of brains from the indicated mice at six weeks of age. An enlarged lateral ventricle with a surrounding edematous area in an *Adar1^W197A/W197A^* mouse are indicated with an asterisk and black arrow, respectively. Scale bars, 1.25 mm. **(E, F)** Representative images of the cerebral cortex of the indicated mice stained with anti-GFAP (**E**) and anti-Iba1 (F) antibodies. Scale bar, 500 μm (**E**) and 200 μm (**F**).

We observed the aberrant development of multiple organs in *Adar1^W197A/W197A^* mice. For instance, the total brain size of these mutant mice was small and accompanied by the thinner cerebral cortical and white matter layers, and a cerebellar granular layer (**Figure 2D**). The lateral and fourth ventricles were enlarged and accompanied by hypoplasia of the choroid plexus. These abnormalities were not observed in the brains of *Adar1^W197A/W197A^ Ifih1^-/-^* mice, which suggests that ADAR1 p150–mediated Z-RNA recognition is required to prevent abnormal brain development in a MDA5-dependent manner. Immunostaining analysis against glial fibrillary acidic protein (GFAP), a marker for astrocytes, demonstrated that the number of astrocytes was increased in the brains of *Adar1^W197A/W197A^* mice, with such cells spreading into the large area, including cerebral cortex and corpus callosum (**Figure 2E**). Astrocytosis has been reported in the brain of a patient with AGS (van Heteren et al., 2008) while astrocytes were thought to be a major source of cytokines that were overproduced in the AGS brain (Cuadrado et al., 2015; Sase et al., 2018). Furthermore, the number of cells positive for ionized calcium binding adapter protein 1 (Iba1), a marker of activated microglia, was also increased in the cerebral cortex (**Figure 2F**). Encephalopathy, with astrocytosis and microgliosis, has been observed in mutant mice with constitutively-activated MDA5, which is encoded by an AGS-causative *Ifih1* gene (Onizawa et al., 2020). However, the pathogenesis of *ADAR1*-mediated encephalopathy found in patients with AGS is poorly understood, given that no animal models manifesting encephalopathy have been established. Therefore, *Adar1^W197A/W197A^* mice will be a valuable resource for understanding the pathogenesis of AGS-associated encephalopathy.

### Hematopoiesis is impaired in *Adar1^W197A/W197A^* mice

We also observed the aberrant development of organs other than the brain in *Adar1^W197A/W197A^* mice. The mucosal layer of the colon was thinner and showed a reduced density of intestinal glands. The thymus and spleen were severely atrophic, and the density and size of white pulp was reduced in the spleen of *Adar1^W197A/W197A^* mice (**Figure 3A, B**). Such abnormalities were not observed in *Adar1^W197A/W197A^ Ifih1^-/-^* mice, suggesting MDA5-dependent mechanisms. In accordance with the atrophic organ morphology, the total numbers of thymocytes and splenocytes in the thymus and spleen, respectively, were significantly reduced in *Adar1^W197A/W197A^* mice (**Figure 3C, D**). Therefore, we examined the development of immune cells. However, we did not observe abnormal thymocyte maturation in the thymi of the mutant mice (**Figure S1A**). In addition, the proportions of major immune subsets, including T cells (CD3^+^), B cells (CD19^+^), granulocytes (CD11b^+^Ly6G^+^), and monocytes (CD11b^+^Ly6G^-^), were not altered in the spleens of *Adar1^W197A/W197A^* mice (**Figure S1B, C**). Furthermore, the proportions of erythroblast subpopulations (R2 to R5) were preserved (**Figure S1D**). Therefore, we next evaluated the transition from lineage marker (L)^-^, c-Kit (K)^+^, Sca-I (S)^+^ cells (LKS^+^ cells) to LKS^-^ cells during hematopoiesis in the bone marrow, which requires ADAR1 expression (Hartner et al., 2009). We observed a significant increase in the population of LKS^+^ cells in *Adar1^W197A/W197A^* mice (**Figure 3E**). These data collectively suggest that the reduction in the number of thymocytes and splenocytes found in these mutant mice is most likely attributed to an impairment of early hematopoiesis, which requires ADAR1 p150–mediated Z-RNA recognition.

**Figure 3.**
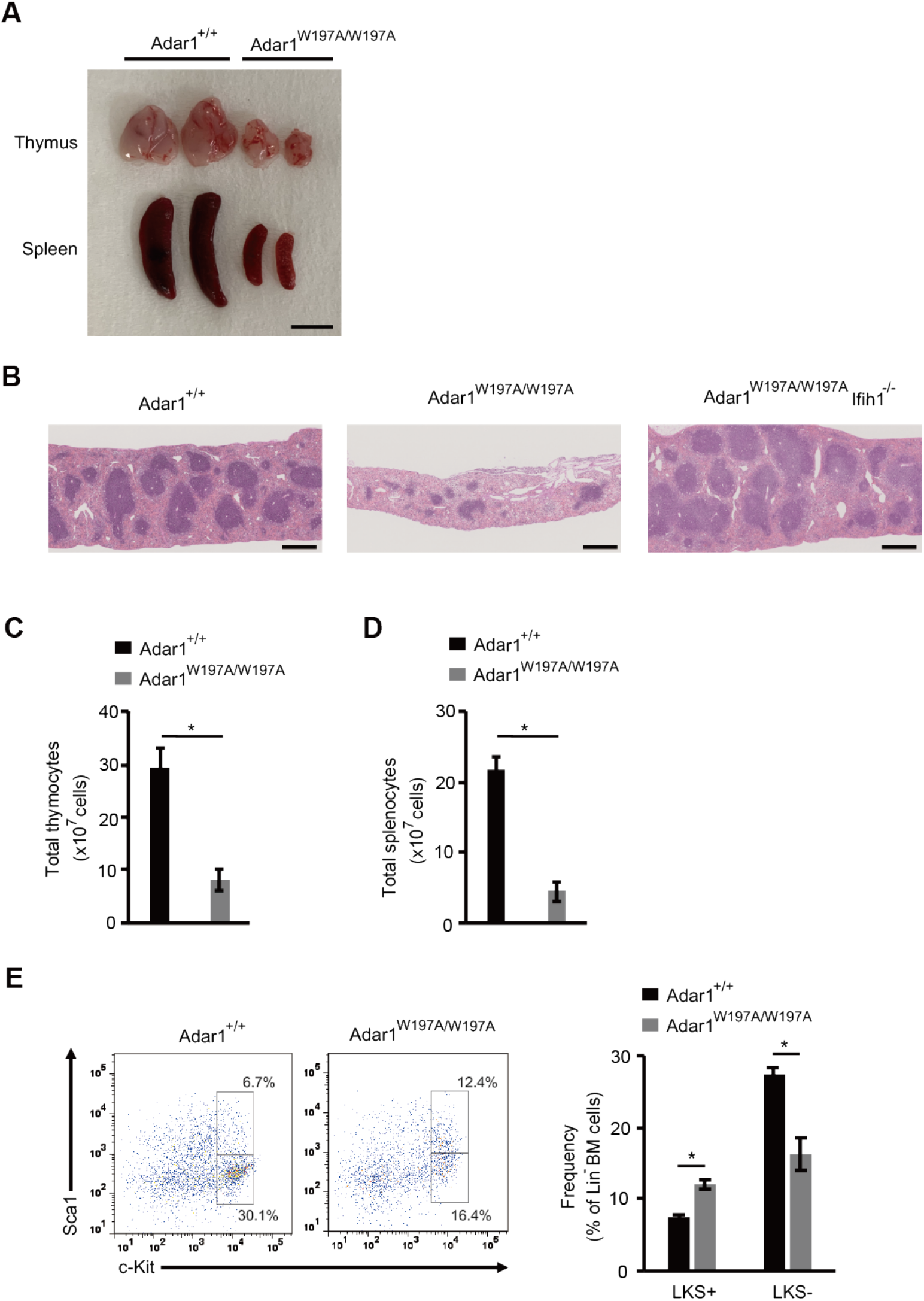
Aberrant hematopoietic stem cell differentiation observed in *Adar1^W197A/W197A^* mice. **(A)** Representative images of thymi and spleens isolated from *Adar1*^+/+^ and *Adar1^W197A/W197A^* mice. Scale bar, 0.5 cm. (**B**) Representative images of hematoxylin and eosin (HE) staining of the spleen at six weeks of age from the indicated mice. Scale bar, 500 μm. (**C, D**) The total number of cells in the thymus (**C**) and spleen (**D**) in *Adar1^+/+^* (n = 4) and *Adar1^W197A/W197A^* (n = 4) mice. Values represent the mean ± SEM (Mann–Whitney *U*-test, \**p* < 0.05). (**E**) Flow cytometric analysis of c-Kit and Sca1 expression in lineage marker-negative bone marrow (BM) cells. Representative flow cytometric images are shown (*left panel*). The percentage of c-Kit^+^Sca1^+^ (LKS+) and c-Kit^+^Sca1^-^ (LKS-) BM cells is compared between *Adar1^+/+^* (n = 4) and *Adar1^W197A/W197A^* (n = 4) mice (*right panel*). Values represent the mean ± SEM (Mann–Whitney *U*-test, \**p* < 0.05).

### Z-RNA–recognition is required for efficient RNA editing and prevention of MDA5 activation *in vivo*

We examined the expression of ISGs in various organs of adult *Adar1^W197A/W197A^* mice. This analysis demonstrated that the expression levels of three ISGs, *Ifit1, Rsad2*, and *Cxcl10*, were significantly upregulated in the mutant mice to various degrees in an organ-dependent manner (**Figure 4A**). Of note, the upregulated expression of ISGs was the most enhanced in the brain, which might be the reason why this is the most affected organ in patients with AGS (Crow et al., 2015). Furthermore, the upregulated expression of ISGs was detected in brains collected at P0 from *Adar1^W197A/W197A^* mice (**Figure 4B**), indicating perinatal onset as reported in patients with AGS (Rice et al., 2007). As expected, concurrent deletion of MDA5 ameliorated the aberrant expression of ISGs, which reconfirmed that the abnormal development and type I IFN signature found in *Adar1^W197A/W197A^* mice are mediated by MDA5 activation (**Figure 4A**).

**Figure 4.**
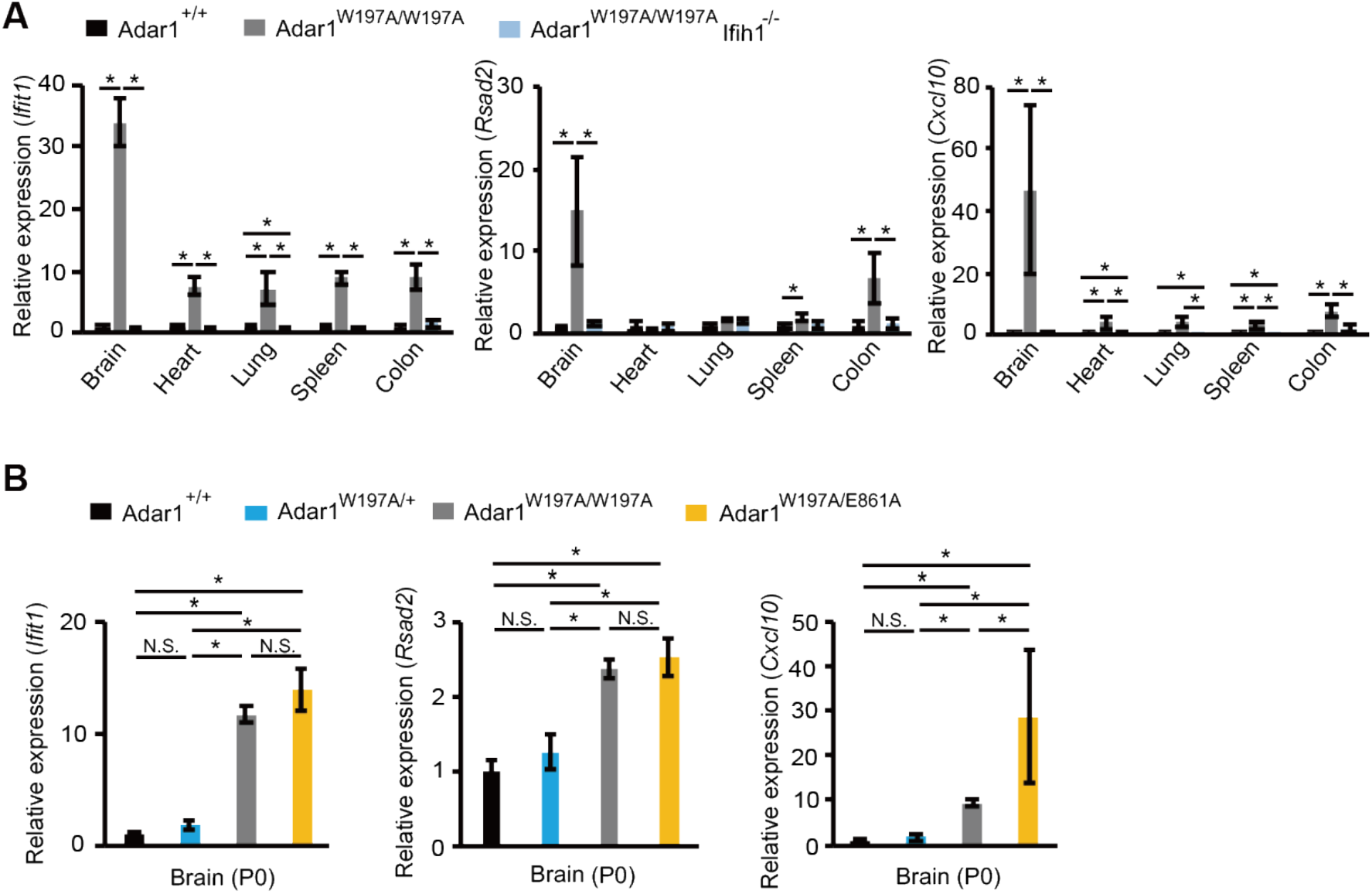
The expression of ISGs is upregulated in *Adar1^W197A/W197A^* mice. **(A)** The relative expression of the mRNAs of the type I interferon-stimulated genes (ISGs), *Ifit1, Rsad2, and Cxcl10*, in various organs at six weeks of age is compared between *Adar1^+/+^* (n = 3), *Adar1^W197A/W197A^* (n = 3) and *Adar1^W197A/W197A^ Ifih1^-/-^* (n = 3) mice. Values represent relative gene expression normalized to GAPDH mRNA and are displayed as the mean ± SEM (Mann–Whitney *U*-test, \**p* < 0.05). We indicated only significant differences. **(B)** The relative expression of the mRNAs of the ISGs in the brain at P0 is compared among *Adar1^+/+^* (n = 6), *Adar1 ^W197A/+^* (n = 3), *Adar1^W197A/W197A^* (n = 4), and *Adar1^W197A/E861A^* (n = 3) mice. Values represent relative gene expression normalized to GAPDH mRNA and are displayed as the mean ± SEM (Mann–Whitney *U*-test, \**p* < 0.05, N.S., not significant).

Next, we investigated the possibility that Z-RNA–recognition of ADAR1 p150 might contribute to preventing MDA5 activation in an RNA editing–independent manner. For this purpose, we crossed *Adar1^W197A/+^* mice with *Adar1*^E861A/+^ mice because ADAR1 p150 (E861A) loses RNA editing activity but contains an intact Zα domain. However, the upregulated expression of ISGs was not ameliorated in the brains of *Adar1^W197A/E861A^* mice, although the expression of ISGs in the brains of *Adar1^W197A/+^* mice was not significantly different from that in *Adar1^+/+^* mice (**Figure 4B**). Collectively, these data suggest that ADAR1 p150–mediated recognition of Z-RNA is required for RNA editing of certain sites, which is essential for avoiding MDA5 activation.

To identify RNA editing sites affected by Z-RNA recognition of ADAR1 p150 *in vivo*, we examined the expression level of Adar1 p150 mRNA in *Adar1^W197A/W197A^* mice. This analysis demonstrated that the expression level of Adar1 p150 mRNA was significantly upregulated ~2-to-3–fold in the brains and hearts of such mutant mice (**Figure 5A**). We have previously reported that the expression of ADAR1 p150 is extremely low in the mouse brain where ADAR1 p110 is abundantly expressed (Nakahama et al., 2018). However, ADAR1 p150 was detectable in the brains of *Adar1^W197A/W197A^* mice, which was in accordance with the upregulated expression of Adar1 p150 mRNA (**Figure 5B**). In addition, an increased amount of ADAR1 p150 protein was also observed in the hearts of these mutant mice. Given that Adar1 p150 transcription is triggered by an IFN-inducible promoter (George et al., 2005; George et al., 2008; George and Samuel, 1999), these results suggest that the expression of ADAR1 p150 was increased in a feedback manner to compensate for reduced RNA editing at certain ADAR1 p150–specific sites that require Z-RNA recognition for efficient editing. Therefore, we performed poly(A) RNA-seq and compared the editing ratio of each RNA site in the brains of *Adar1^+/+^* and *Adar1 ^W197A/W197A^* mice. In accordance with the increased expression of ADAR1 p150, we observed that the editing ratio of ~30% of sites in *Adar1^W197A/W197A^* mice was more than 10% higher than that in *Adar1^+/+^* mice (**Figure 5C, Supplementary Table S2**). Zα domain–mediated Z-RNA recognition might not be essential for RNA editing of these sites, which might also be editable by ADAR1 p110. ADAR1 p110 is abundantly expressed in the brain and seemed to be slightly increased in a feedback manner in the brains of *Adar1^W197A/W197A^* mice (**Figure 5B**) as previously reported (Steinman and Wang, 2011). However, this compensatory upregulation of RNA editing could not ameliorate an inflammatory ISG signature (**Figure 4A, B**), indicating that certain sites require Z-RNA-binding for efficient editing, which is indispensable for preventing MDA5 activation. Indeed, the editing ratio of ~5% of sites in *Adar1^W197A/W197A^* mice was more than 10% lower than that in *Adar1^+/+^* mice even under the condition of increased expression of ADAR1 p150 (**Figure 5C, Supplementary Table S2**). Of these sites, the editing ratios of a very limited number of the sites were commonly reduced in dKO cells expressing ADAR1 p150 (W197A) (**Figure 5D**). These sites are top candidates that require Z-RNA recognition for their efficient editing and which needs further investigation to specify. Finally, we observed that the increased expression of ADAR1 p150 and the resultant upregulation of RNA editing was ameliorated in *Adar1^W197A/W197A^ Ifih1^-/-^* mice (**Figure 5A–D**). Collectively, these results suggest that although the number of sites may be very limited, at least in the brain, Z-RNA–mediated RNA editing of certain sites is required for preventing MDA5 activation.

**Figure 5.**
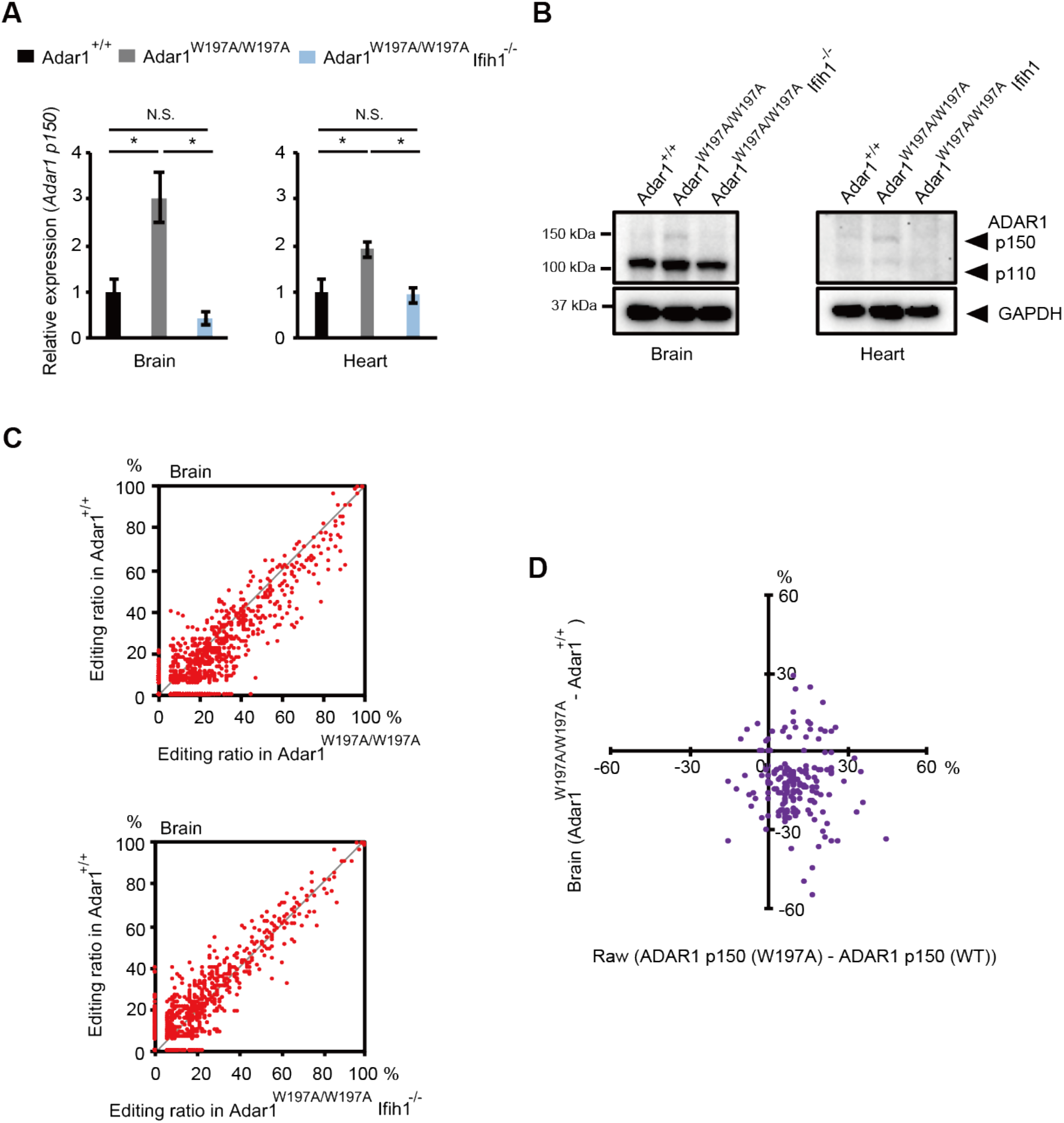
RNA editing is upregulated in a feedback manner in *Adar1^W197A/W197A^* mice. **(A)** The relative expression of Adar1 p150 mRNA in the brain and heart at six weeks of age is compared between *Adar1^+/+^* (n = 3), *Adar1^W197A/W197A^* (n = 3), and *Adar1^W197A/W197A^ Ifih1^-/-^* (n = 3) mice. Values represent relative gene expression normalized to GAPDH mRNA and are displayed as the mean ± SEM (Mann–Whitney *U*-test, *\*p* < 0.05, N.S., not significant). (**B**) Immunoblot analysis of ADAR1 p110 and p150 protein expression in brains and hearts isolated from indicated mice at six weeks of age. The expression of GAPDH protein is shown as a reference. (**C**) The mean editing ratio in the brain isolated at three weeks of age (n = 2 for each group) is compared between *Adar1^+/+^* and *Adar1^W197A/W197A^* mice (*upper panel*), and *Adar1^+/+^* and *Adar1^W197A/W197A^ Ifih1^-/-^* mice (*lower panel*). We considered only the sites that are covered with more than 20 reads in all samples. **(D)** The mean editing ratio of each site in the brain of *Adar1*^+/+^ mice (n = 2) was subtracted from that of *Adar1^W197A/W197A^* mice (n = 2) and the resultant absolute value was plotted on the horizontal axis. The mean editing ratio of each site in *Adar1/Adar2* dKO Raw 264.7 cells expressing wild-type (WT) ADAR1 p150 (n = 2) was subtracted from that in the cells expressing Zα domain–mutated (W197A) ADAR1 p150 and the resultant absolute value was plotted on the vertical axis. We considered only the sites that were commonly detected in mice (covered with more than 20 reads) and cultured cells (covered with more than 5 reads).

In summary, we elucidated that Z-RNA–binding of ADAR1 p150 is necessary for maintaining the editing ratio of certain sites at a proper level. In addition, we found that disrupted Z-RNA–binding of ADAR1 p150 results in activation of the MDA5 pathway in an RNA editing–dependent manner. By inserting a point mutation in the Zα domain to abolish Z-RNA–binding, we successfully established *Adar1* mutant mice that manifest abnormal organ development with a type I IFN signature by avoiding embryonic lethality for the first time. This includes encephalopathy accompanied by gliosis, which is reminiscent of the phenotype found in patients with AGS. These findings in mice and cell culture models shed light on the biological significance of Z-RNA, especially for regulating the immune system.

## MATERIALS AND METHODS

### Mice maintenance

Mice were maintained on a 12-h light/dark cycle at a temperature of 23 ± 1.5°C with humidity of 45 ± 15% as previously described (Vongpipatana et al., 2020). All experimental procedures that included mice were performed following protocols approved by the Institutional Animal Care and Use Committee of Osaka University.

### Mutant mice

*Ifih1^-/-^* and *Adar1^E861A/E861A^* mice were maintained in our laboratory as previously described (Costa Cruz et al., 2020). *Adar1^W197A/W197A^* and *Adar1^N175A·W197A/ N175A·W197A^* mice were generated by genome editing using a CRISPR/Cas9 system at the Genome Editing Research and Development Center, Graduate School of Medicine, Osaka University. Briefly, Alt-R CRISPR-Cas9 CRISPR RNA (5’-AAGGGCACAAGGCTCCACAA-3’) was synthesized at Integrated DNA Technologies (IDT) and hybridized with trans-activating CRISPR RNA (tracrRNA), generating guide RNAs. Pronuclear-stage mouse embryos obtained from *Ifih1^-/-^* mice were electroporated to introduce Cas9 mRNA, the guide RNA, and a single-stranded donor (5’-TGTGCTAGCCAGAGAGCTCAGAATCCCCAAAAGGGACAT<u>GCC</u>CGTATTTTG TACTCCCTGGAAAAGAAGGGAAAGCTGCACAGAGGAAGGGGGAAACCTCC TTTG<u>GCC</u>AGCCTTGTGCCCTTGAGTCAGGCTTGGACTCAGCCCCCTGG-3’); these introduced either a single or double point mutation(s) at the corresponding codons (underlined GCC in the target nucleotide). Mouse embryos that developed to the two-cell stage were transferred into the oviducts of female surrogates.

For genotyping of *Adar1^W195A/W195A^* and *Adar1^N173A·W195A/ N173A·W195A^* mice, genomic DNA was amplified with the following primers: 5’-CTCAGAAATCAGGGGTTCCA-3’ and 5’-CTTGGTGAGGCCAATGTTCT-3’. After treating with illustra ExoProStar (GE Healthcare), PCR products were directly sequenced using the primer 5’-ATAGGACCCCCACTTCCTCC-3’. All mice used in experiments were in a C57BL/6J background.

### Cell culture

Mouse macrophage-like Raw 264.7 cells were cultured in Dulbecco’s modified Eagle’s medium (Nacalai Tesque) supplemented with 10% fetal bovine serum and 1% penicillin/streptomycin (Thermo Fisher Scientific). Cells were maintained at 37°C in the presence of 5% CO_2_.

### Generation of *Adar1/Adar2* double knockout Raw 264.7 cells

*Adar1/Adar2* dKO Raw 264.7 cells were generated by genome editing using a CRISPR/Cas9 system. In brief, a pSpCas9 BB-2A-GFP (PX458) plasmid that contained predesigned guide RNA targeting *Adar1* (5’-CTTGTCCGTCAAGTACCAGA-3’) and a pLentiCRISPR v2 plasmid that contained predesigned guide RNA targeting *Adar2* (5’-AGTACCGCCTGAAGAAGCGA-3’) were obtained from GenScript. These plasmids were then transfected into Raw 264.7 cells using a Neon Transfection System (Thermo Fisher Scientific) with the following parameters: pulse voltage, 1,680 V; pulse width, 20 ms; and pulse number, 1. Twenty-four hours after transfection, GFP-positive cells were sorted using an SH800 cell sorter (Sony). After clonal expansion of each cell, genomic DNA was extracted and regions encompassing *Adar1* and *Adar2* genes were amplified by PCR with the following primers: 5’-CATGGCTGAAATCAAGGAGAAGATC-3’ and 5’-TCATGCCTACTTTCATGCTTTATCG-3’ for *Adar1*, and 5’-GTTGTAAGTTACTCTTTCTGGGCAC-3’ and 5’-GTTTACCTCCACAGACATGACAAAC-3’ for *Adar2*. After treating with illustra ExoProStar (GE Healthcare), PCR products were directly sequenced using the following primers: 5’-TGGCTGAAATCAAGGAGAAGATCT-3’ for *Adar1*, and 5’-ACAGGTTCCAGCAGCACTG-3’ for *Adar2* to screen *Adar1*/*Adar2* dKO cells.

### Construction of plasmids

A plasmid containing mouse ADAR1 p150 was obtained from DNAFORM (Yokohama, Japan). The coding region of ADAR1 p150 was amplified by PCR and the resultant PCR product was inserted into a pEGFP-C1 vector (Clontech) using XhoI/BamHI restriction enzyme–recognition sites to generate the expression construct, which was termed pEGFP-C1-mADAR1 p150 (WT). To insert a W195A point mutation in the Zα domain, site-directed mutagenesis was performed using a PrimeSTAR mutagenesis basal kit (Takara Bio) with the following primers: 5’-TCCTTTGGCGAGCCTTGTGCCCTTGAG-3’ and 5’-AAGGCTCGCCAAAGGAGGTTTCCCCCT-3’. The resultant expression construct was termed pEGFP-C1-mADAR1 p150 (W195A). Similarly, to insert triple mutations in the three dsRBDs (dsRBD1; KKVAK to EAVAA, dsRBD2; KKVAK to EAVAA and dsRBD3; KKQGK to EAQGA) to abolish binding capacity to dsRNA (Valente and Nishikura, 2007), site-directed mutagenesis was performed using a PrimeSTAR mutagenesis basal kit (Takara Bio) with the following primers: 5’-TGGCAGCGAGGCAGTAGCCGCGCAGGACGCAGCAGTGAA-3’ and 5’-GTCCTGCGCGGCTACTGCCTCGCTGCCAGCCTCAGCTGG-3’ for site1, and 5’-CCCCAGCGAGGCGGTAGCAGCGCAGATGGCCGCAGAGGA-3’ and 5’-CATCTGCGCTGCTACCGCCTCGCTGGGGGCGCTCACAGG-3’ for site 2, and 5’-ACACAGCGAGGCACAGGGCGCGCAGGATGCAGCGGATGC-3’ and 5’-ATCCTGCGCGCCCTGTGCCTCGCTGTGTGCACACACGGC-3’ for site 3. The resultant expression construct was termed pEGFP-C1-mADAR1 p150 (mut_dsRBDs). All the constructs were verified by DNA sequencing.

### Plasmid transfection

Plasmids pEGFP-C1-mADAR1 p150 (WT), pEGFP-C1-mADAR1 p150 (W195A), or pEGFP-C1-mADAR1 p150 (mut_dsRBDs) was transfected into *Adar1/Adar2* dKO Raw 264.7 cells using a Neon Transfection System (Thermo Fisher Scientific) with the following parameters: pulse voltage, 1,750 V; pulse width, 25 ms; and pulse number, 1. Twenty-four hours after transfection, GFP-positive cells were sorted using an SH800 cell sorter and subjected to RNA extraction.

### Total RNA extraction

Total RNA was extracted from isolated organs or collected cells using TRIzol reagent (Thermo Fisher Scientific) by following the manufacturer’s instructions. After measuring the RNA concentration using a NanoDrop One (Thermo Fisher Scientific), total RNA samples were stored at −80°C until use.

### Poly(A) RNA sequencing analysis

Poly(A) RNA sequencing (RNA-seq) analyses were performed at Macrogen (Kyoto, Japan) from a library preparation using a TruSeq Stranded mRNA Library Prep kit (Illumina). The library samples were subjected to deep sequencing using Illumina NovaSeq 6000 sequencing platforms with 101-bp paired-end reads.

### Genome-wide identification of RNA editing sites

We adopted a genome-wide approach to identify editing sites within RNA-seq reads as previously described (Costa Cruz et al., 2020; Nakahama et al., 2018) but with modifications, based in part on the literature (Liddicoat et al., 2015; Ramaswami et al., 2013). In brief, sequence reads were mapped onto a reference mouse genome (GRCm38/mm10) with a spliced aligner HISAT2 (Kim et al., 2015). The mapped reads were then processed by adding read groups, and sorting and marking duplicates with the tools, AddOrReplaceReadGroups and MarkDuplicates, compiled in GATK4 (McKenna et al., 2010). GATK SplitNCigarReads, BaseRecalibrator, and ApplyBQSR were used to split ‘N’ trim and reassign mapping qualities, which output analysis-ready reads for subsequent variant calling. The GATK HaplotypeCaller was run for variant detection, in which the stand-call-conf option was set to 20.0 and the dont-use-soft-clipped-bases option was used. The results of variant calling were further filtered by GATK VariantFiltration using Fisher strand (FS) values > 30.0 and quality by depth (QD) values < 2.0 as recommended by the GATK developer for RNA-seq analysis. The remaining variants that were expected to be of high quality were annotated with ANNOVAR software (Wang et al., 2010), where gene-based annotation was generated with a RefSeq database (O’Leary et al., 2016). Among these variants, we picked up known editing sites registered in the following databases: database of RNA editing (DARNED) (Kiran et al., 2013), rigorously annotated database of A-to-I RNA editing (RADAR) (Ramaswami and Li, 2014), or REDIportal (Lo Giudice et al., 2020). Of note, DARNED and REDIportal provide editing sites in mm10 coordinates, while RADAR does in mm9. This discordance was resolved in a way that RADAR-registered editing sites were uplifted to mm10 coordinates with a liftOver tool in University of California Santa Cruz (UCSC) Genome Browser utilities (Lee et al., 2020). It should also be noted that the strand orientation in the predicted variants was strictly checked with that of the database annotations. Finally, A-to-I editing ratios in each sample were calculated by dividing the allelic depth by the read depth for the editing sites shown in the annotated results.

### qRT–PCR analysis

As previously described (Nakahama et al., 2018; Vongpipatana et al., 2020), cDNA was synthesized from total RNA extracted from various organs using a ReverTra Ace qPCR–RT Master Mix with guide DNA Remover (Toyobo). The quantitative reverse transcription (qRT)–PCR reaction mixture was prepared by combining each target-specific primers and probes with a THUNDERBIRD Probe qPCR Mix (Toyobo). The qRT–PCR was performed using an ABI Prism 7900HT Real-Time PCR System (Applied Biosystems). The sequences of primers and probes for *Adar1 p150, Ifit1, Cxcl10, Rsad2*, and *GAPDH* have been previously reported (Nakahama et al., 2018). The expression level of each mRNA relative to that of GAPDH mRNA was calculated by the ΔΔCt method.

### Immunoblot analysis

Tissue and cell lysates were prepared as described previously with some modifications (Miyake et al., 2016). In brief, isolated organs were frozen in liquid nitrogen, thawed once at room temperature, and then homogenized in lysis buffer (0.175 M Tris-HCl, pH 6.8, 3% SDS, and 5 mM EDTA). After boiling at 95°C for 10 min, the lysates were passed through a 23-gauge needle followed by centrifugation at 20,000 × g and 4°C for 10 min. Each supernatant was transferred to a 1.5 mL tube and stored at −80°C until use. Immunoblot analysis was performed as previously described (Costa Cruz et al., 2020; Nakahama et al., 2018) with minor modifications. In brief, 30 μg of each protein lysate sample was separated by sodium dodecyl sulfate–polyacrylamide gel electrophoresis, transferred to a polyvinylidene difluoride membrane (Bio-Rad), and immunoblotted at 4°C overnight with primary antibodies: mouse monoclonal anti-ADAR1 antibody (15.8.6; Santa Cruz Biotechnology), mouse monoclonal anti-ADAR2 antibody (1.3.1; Santa Cruz Biotechnology), and mouse monoclonal anti-GAPDH (M171-3; MBL).

### Histological analysis

Mice were anesthetized and transcardially perfused with phosphate buffered saline (PBS) followed by 4% paraformaldehyde (PFA) in PBS. Organs were dissected, postfixed in 4% PFA/PBS with shaking at 4°C for 2days, and embedded in paraffin blocks. Hematoxylin and eosin staining, and morphological analysis by pathologists were performed at Morphotechnology Co. Ltd. (Hokkaido, Japan) and Applied Medical Research Laboratory Co. Ltd. (Osaka, Japan).

### Immunohistochemistry

Fixed paraffin-embedded tissues were sectioned at an 8-μm thickness using a microtome (RM2145; Leica). The sections placed onto glass slides were deparaffinized, rehydrated, and treated with antigen retrieval buffer, HistoVT-One (Nacalai Tesque, Japan) at 70°C for 20 min. The sections were then washed three times in 0.01% PBST (0.01% Tween-20 in PBS) for 5 min and incubated with 3% hydrogen peroxide/PBST solution for 20 min, after which they were subsequently washed threee times in PBST for 5 min. The sections were then incubated with primary antibodies dissolved in blocking buffer (0.4% Block ACE [UKB80; KAC, Japan]/PBST) at 4°C overnight, and then washed three times in PBST before incubation with biotinylated-secondary antibodies dissolved in PBST (BA-1000 for anti-rabbit and BA-9200 for anti-mouse; Vector Laboratories, 1:1,000 dilution) at 25°C for 1 h. After another round of washing, sections were treated with Avidin-Biotin Complex (PK-4000, Vector Laboratories) for 1 h at 25°C and washed three times in PBST for 5 min. Sections were then treated with 3,3’-diaminobenzidine (DAB; D4293, Sigma-Aldrich) until the desired brown stain intensity developed, and washed in PBST. Sections were then dehydrated in an ascending series of diluted ethanols, treated with xylene, mounted in xylene-based mounting medium (Entellan new; Merck Millipore) and coverslips placed on top. Primary antibodies used were rabbit polyclonal anti-Iba1 antibody (019-19741; Fujifilm Wako, Japan; 1:1,000), and mouse monoclonal anti-GFAP antibody (G3893; Sigma-Aldrich; 1:1,000). Images were captured using an Olympus BX63 fluorescence microscope.

### Flow cytometry

Freshly isolated thymus or spleen was mashed through a 70-μm cell strainer (Falcon) to yield single-cell suspensions. Bone marrow cells were flushed from tibiae and femurs. Cells were then counted and stained with fluorescent dye–conjugated antibodies against CD3 (17A2; Tonbo Biosciences), CD4 (RM4-5; BioLegend), CD8 (53-6.7; BioLegend), CD11b (M1/70; BioLegend), CD19 (1D3; Tonbo Biosciences), CD71 (RI7217; BioLegend), CD117 (2B8; BioLegend), Ly6G (1A8; BioLegend), ScaI (D7; BioLegend), Ter119 (TER-119; BioLegend), and lineage markers (133301; BioLegend). After incubating cells with antibodies at 4°C for 30 min, the cells were washed with Dulbecco’s PBS and analyzed by using a FACSCanto II flow cytometer (BD) and FlowJo software (Tree Star).

### Statistical analysis

A Mann–Whitney *U*-test was used as indicated in each figure legend. All values are displayed as the mean ± standard error of the mean (SEM). Non-significance is displayed as N.S., while statistical significance is displayed as *p* < 0.05 (*) or *p* < 0.01 (**).

## ACCESSION NUMBERS

The RNA-seq data used in this study are available through the DNA Data Bank of Japan (DDBJ) under accession number DRA011210.

## ACKNOWLEDGMENTS

We thank all staff at the Genome Editing Research and Development Center, the Center for Medical Research and Education, the CoMIT Omics Center, and the Institute of Experimental Animal Sciences, Graduate School of Medicine, Osaka University, for technical support. Computations were partially performed on the NIG supercomputer at ROIS National Institute of Genetics, Japan. This work was supported by Grants-in-Aid KAKENHI (19K22580 and 20H03341 to Y. Kawahara, 18K15186 to T.N., 20J11266 to J.I.K., and 18K11526 to Y. Kato) from the Ministry of Education, Culture, Sports, Science and Technology (MEXT) of Japan, a grant (JP20ek0109433 to T.N.) from the Japan Agency for Medical Research and Development (AMED), and by grants from The Tokyo Biochemical Research Foundation (to Y. Kawahara), The Naito Foundation, and Novartis Research Grants, The Mochida Memorial Foundation for Medical and Pharmaceutical Research, Astellas Foundation for Research on Metabolic Disorders, The Uehara Memorial Foundation, The Osaka Medical Research Foundation for Intractable Diseases (to T.N.), and the Takeda Science Foundation (to Y. Kawahara and T.N.). F.X. was supported by MEXT scholarships. J.I.K. was supported by The Korean Scholarship Foundation. J.I.K. and T.V. were supported by a Research Fellowship for Young Scientists from the Japan Society for the Promotion of Science (JPSP).

## AUTHOR CONTRIBUTIONS

T.N. and Y. Kawahara designed the study and wrote the manuscript. Y. Kato. performed all the bioinformatics analyses. T.N., T.S., J.I.K., T.V., H.T., Y.X., and Y. Kawahara performed all other experiments. All authors agree to the content of the final manuscript.

## DECLARATION OF INTERESTS

The authors declare no competing interests.

## SUPPLEMENTARY FIGURES AND FIGURE LEGENDS

**Figure S1.**
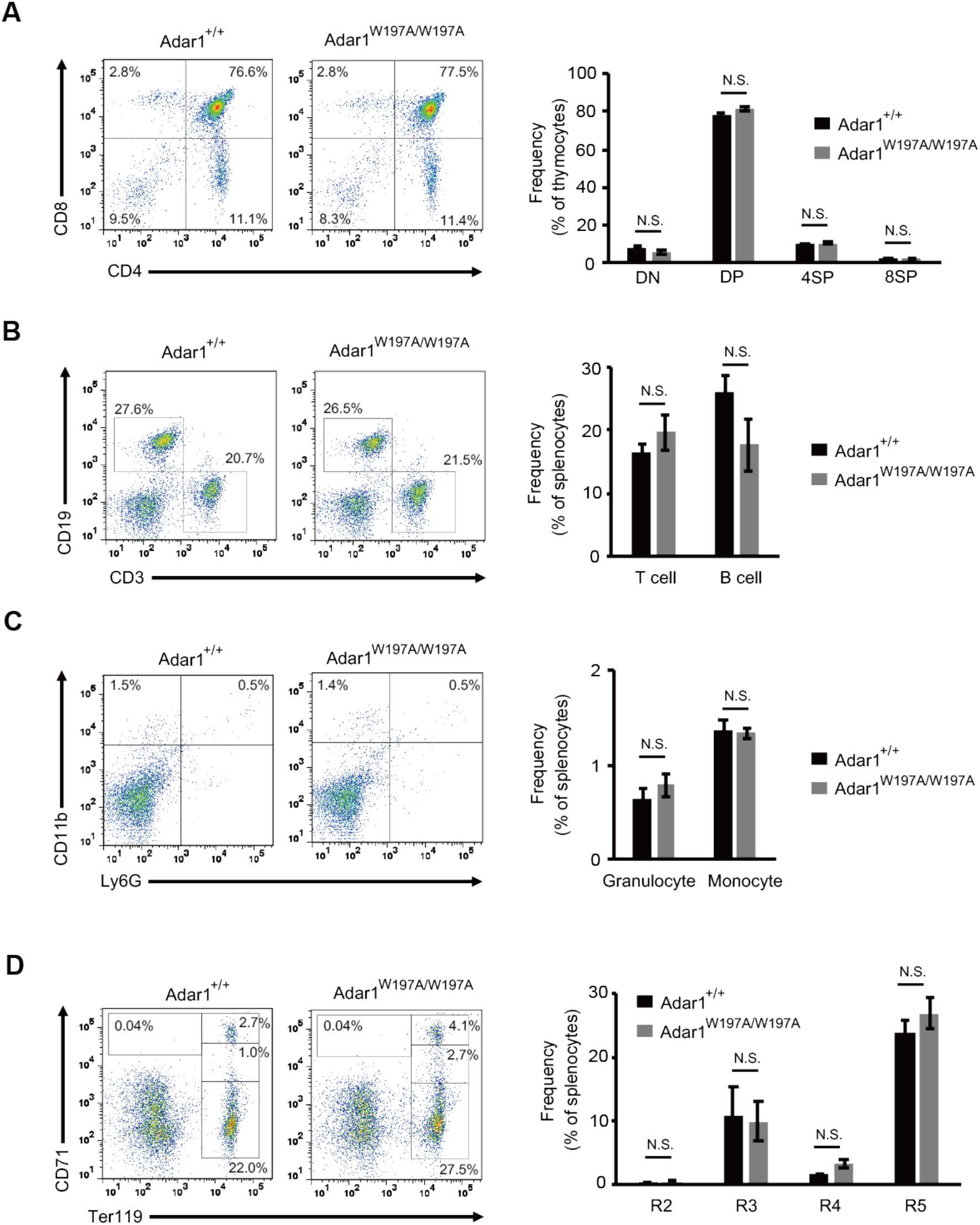
Analysis of the thymocytes and splenocytes in *Adar1^W197A/W197A^* mice. **(A)** Representative flow cytometric images of CD4 and CD8 expression on thymocytes (*left panel*). The percentages of CD4^-^CD8^-^ double-negative (DN), CD4^+^CD8^+^ double-positive (DP), CD4^+^CD8^-^ single-positive (4SP), and CD4^-^CD8^+^ single-positive (8SP) thymocytes are compared between *Adar1^+/+^* (n = 4) and *Adar1^W197A/W197A^* (n = 4) mice (*right panel*). **(B)** Representative flow cytometric images of CD3 and CD19 expression on splenocytes (*left panel*). The percentages of CD3^+^CD19^-^ (T cells) and CD3^-^CD19^+^ (B cells) cells are compared between *Adar1^+/+^* (n = 4) and *Adar1^W197A/W197A^* (n = 4) mice (*right panel*). (**C**) Representative flow cytometric images of Ly6G and CD11b expression on splenocytes (*left panel*). The percentages of Ly6G^+^CD11b^-^ (Granulocyte) and Ly6G^-^CD11b^+^ (Monocyte) cells are compared between *Adar1^+/+^* (n = 4) and *Adar1^W197A/W197A^* (n = 4) mice (*right panel*). (**D**) Representative flow cytometric images of Ter119 and CD71 expression on splenocytes (*left panel*). The percentages of Ter119^-^CD71^hi^ (R2), Ter119^+^CD71^hi^ (R3), Ter119^+^CD71^med^ (R4), and Ter119^+^CD71^lo^ (R5) erythroid progenitor cells are compared between *Adar1^+/+^* (n = 4) and *Adar1^W197A/W197A^* (n = 4) mice (*right panel*). Values represent the mean ± SEM (Mann–Whitney *U*-test, N.S., not significant).

## Notes

### Competing Interest Statement

The authors have declared no competing interest.

## REFERENCES

Bass, B.L. (2002). RNA editing by adenosine deaminases that act on RNA. Annu Rev Biochem 71, 817–846.

Bazak, L., Haviv, A., Barak, M., Jacob-Hirsch, J., Deng, P., Zhang, R., Isaacs, F.J., Rechavi, G., Li, J.B., Eisenberg, E., and Levanon, E.Y. (2014). A-to-I RNA editing occurs at over a hundred million genomic sites, located in a majority of human genes. Genome Res 24, 365–376.

Brown, B.A., 2nd, Lowenhaupt, K., Wilbert, C.M., Hanlon, E.B., and Rich, A. (2000). The zalpha domain of the editing enzyme dsRNA adenosine deaminase binds lefthanded Z-RNA as well as Z-DNA. Proc Natl Acad Sci U S A 97, 13532–13536.

Costa Cruz, P.H., Kato, Y., Nakahama, T., Shibuya, T., and Kawahara, Y. (2020). A comparative analysis of ADAR mutant mice reveals site-specific regulation of RNA editing. RNA 26, 454–469.

Crow, Y.J., Chase, D.S., Lowenstein Schmidt, J., Szynkiewicz, M., Forte, G.M., Gornall, H.L., Oojageer, A., Anderson, B., Pizzino, A., Helman, G., et al. (2015). Characterization of human disease phenotypes associated with mutations in TREX1, RNASEH2A, RNASEH2B, RNASEH2C, SAMHD1, ADAR, and IFIH1. Am J Med Genet A 167A, 296–312.

Cuadrado, E., Michailidou, I., van Bodegraven, E.J., Jansen, M.H., Sluijs, J.A., Geerts, D., Couraud, P.O., De Filippis, L., Vescovi, A.L., Kuijpers, T.W., and Hol, E.M. (2015). Phenotypic variation in Aicardi-Goutieres syndrome explained by cell-specific IFN-stimulated gene response and cytokine release. J Immunol 194, 3623–3633.

George, C.X., Das, S., and Samuel, C.E. (2008). Organization of the mouse RNA-specific adenosine deaminase Adar1 gene 5’-region and demonstration of STAT1-independent, STAT2-dependent transcriptional activation by interferon. Virology 380, 338–343.

George, C.X., and Samuel, C.E. (1999). Human RNA-specific adenosine deaminase ADAR1 transcripts possess alternative exon 1 structures that initiate from different promoters, one constitutively active and the other interferon inducible. Proc Natl Acad Sci U S A 96, 4621–4626.

George, C.X., Wagner, M.V., and Samuel, C.E. (2005). Expression of interferon-inducible RNA adenosine deaminase ADAR1 during pathogen infection and mouse embryo development involves tissue-selective promoter utilization and alternative splicing. J Biol Chem 280, 15020–15028.

Hartner, J.C., Schmittwolf, C., Kispert, A., Muller, A.M., Higuchi, M., and Seeburg, P.H. (2004). Liver disintegration in the mouse embryo caused by deficiency in the RNA-editing enzyme ADAR1. J Biol Chem 279, 4894–4902.

Hartner, J.C., Walkley, C.R., Lu, J., and Orkin, S.H. (2009). ADAR1 is essential for the maintenance of hematopoiesis and suppression of interferon signaling. Nat Immunol 10, 109–115.

Heraud-Farlow, J.E., and Walkley, C.R. (2020). What do editors do? Understanding the physiological functions of A-to-I RNA editing by adenosine deaminase acting on RNAs. Open Biol 10, 200085.

Herbert, A. (2019). Z-DNA and Z-RNA in human disease. Commun Biol 2, 7.

Herbert, A., Alfken, J., Kim, Y.G., Mian, I.S., Nishikura, K., and Rich, A. (1997). A Z-DNA binding domain present in the human editing enzyme, double-stranded RNA adenosine deaminase. Proc Natl Acad Sci U S A 94, 8421–8426.

Herbert, A., Lowenhaupt, K., Spitzner, J., and Rich, A. (1995). Chicken doublestranded RNA adenosine deaminase has apparent specificity for Z-DNA. Proc Natl Acad Sci U S A 92, 7550–7554.

Hsiao, Y.E., Bahn, J.H., Yang, Y., Lin, X., Tran, S., Yang, E.W., Quinones-Valdez, G., and Xiao, X. (2018). RNA editing in nascent RNA affects pre-mRNA splicing. Genome Res 28, 812–823.

Huntley, M.A., Lou, M., Goldstein, L.D., Lawrence, M., Dijkgraaf, G.J., Kaminker, J.S., and Gentleman, R. (2016). Complex regulation of ADAR-mediated RNA-editing across tissues. BMC Genomics 17, 61.

Kawakubo, K., and Samuel, C.E. (2000). Human RNA-specific adenosine deaminase (ADAR1) gene specifies transcripts that initiate from a constitutively active alternative promoter. Gene 258, 165–172.

Kim, D., Langmead, B., and Salzberg, S.L. (2015). HISAT: a fast spliced aligner with low memory requirements. Nat Methods 12, 357–360.

Kiran, A.M., O’Mahony, J.J., Sanjeev, K., and Baranov, P.V. (2013). Darned in 2013: inclusion of model organisms and linking with Wikipedia. Nucleic Acids Res 41, D258–261.

Koeris, M., Funke, L., Shrestha, J., Rich, A., and Maas, S. (2005). Modulation of ADAR1 editing activity by Z-RNA in vitro. Nucleic Acids Res 33, 5362–5370.

Lee, A.R., Kim, N.H., Seo, Y.J., Choi, S.R., and Lee, J.H. (2018). Thermodynamic Model for B-Z Transition of DNA Induced by Z-DNA Binding Proteins. Molecules 23.

Lee, C.M., Barber, G.P., Casper, J., Clawson, H., Diekhans, M., Gonzalez, J.N., Hinrichs, A.S., Lee, B.T., Nassar, L.R., Powell, C.C., et al. (2020). UCSC Genome Browser enters 20th year. Nucleic Acids Res 48, D756–D761.

Levanon, E.Y., Eisenberg, E., Yelin, R., Nemzer, S., Hallegger, M., Shemesh, R., Fligelman, Z.Y., Shoshan, A., Pollock, S.R., Sztybel, D., et al. (2004). Systematic identification of abundant A-to-I editing sites in the human transcriptome. Nat Biotechnol 22, 1001–1005.

Licht, K., Kapoor, U., Amman, F., Picardi, E., Martin, D., Bajad, P., and Jantsch, M.F. (2019). A high resolution A-to-I editing map in the mouse identifies editing events controlled by pre-mRNA splicing. Genome Res 29, 1453–1463.

Liddicoat, B.J., Piskol, R., Chalk, A.M., Ramaswami, G., Higuchi, M., Hartner, J.C., Li, J.B., Seeburg, P.H., and Walkley, C.R. (2015). RNA editing by ADAR1 prevents MDA5 sensing of endogenous dsRNA as nonself. Science 349, 1115–1120.

Lo Giudice, C., Tangaro, M.A., Pesole, G., and Picardi, E. (2020). Investigating RNA editing in deep transcriptome datasets with REDItools and REDIportal. Nat Protoc 15, 1098–1131.

Mannion, N.M., Greenwood, S.M., Young, R., Cox, S., Brindle, J., Read, D., Nellaker, C., Vesely, C., Ponting, C.P., McLaughlin, P.J., et al. (2014). The RNA-editing enzyme ADAR1 controls innate immune responses to RNA. Cell Rep 9, 1482–1494.

McKenna, A., Hanna, M., Banks, E., Sivachenko, A., Cibulskis, K., Kernytsky, A., Garimella, K., Altshuler, D., Gabriel, S., Daly, M., and DePristo, M.A. (2010). The Genome Analysis Toolkit: a MapReduce framework for analyzing next-generation DNA sequencing data. Genome Res 20, 1297–1303.

Miyake, K., Ohta, T., Nakayama, H., Doe, N., Terao, Y., Oiki, E., Nagatomo, I., Yamashita, Y., Abe, T., Nishikura, K., et al. (2016). CAPS1 RNA Editing Promotes Dense Core Vesicle Exocytosis. Cell Rep 17, 2004–2014.

Nakahama, T., Kato, Y., Kim, J.I., Vongpipatana, T., Suzuki, Y., Walkley, C.R., and Kawahara, Y. (2018). ADAR1-mediated RNA editing is required for thymic self-tolerance and inhibition of autoimmunity. EMBO Rep 19.

Nishikura, K. (2010). Functions and regulation of RNA editing by ADAR deaminases. Annu Rev Biochem 79, 321–349.

Nishikura, K. (2016). A-to-I editing of coding and non-coding RNAs by ADARs. Nat Rev Mol Cell Biol 17, 83–96.

O’Leary, N.A., Wright, M.W., Brister, J.R., Ciufo, S., Haddad, D., McVeigh, R., Rajput, B., Robbertse, B., Smith-White, B., Ako-Adjei, D., et al. (2016). Reference sequence (RefSeq) database at NCBI: current status, taxonomic expansion, and functional annotation. Nucleic Acids Res 44, D733–745.

Onizawa, H., Kato, H., Kimura, H., Kudo, T., Soda, N., Shimizu, S., Funabiki, M., Yagi, Y., Nakamoto, Y., Priller, J., et al. (2020). Aicardi-Goutieres syndrome-like encephalitis in mutant mice with constitutively active MDA5. Int Immunol.

Patterson, J.B., and Samuel, C.E. (1995). Expression and regulation by interferon of a double-stranded-RNA-specific adenosine deaminase from human cells: evidence for two forms of the deaminase. Mol Cell Biol 15, 5376–5388.

Pestal, K., Funk, C.C., Snyder, J.M., Price, N.D., Treuting, P.M., and Stetson, D.B. (2015). Isoforms of RNA-Editing Enzyme ADAR1 Independently Control Nucleic Acid Sensor MDA5-Driven Autoimmunity and Multi-organ Development. Immunity 43, 933–944.

Placido, D., Brown, B.A., 2nd, Lowenhaupt, K., Rich, A., and Athanasiadis, A. (2007). A left-handed RNA double helix bound by the Z alpha domain of the RNA-editing enzyme ADAR1. Structure 15, 395–404.

Ramaswami, G., and Li, J.B. (2014). RADAR: a rigorously annotated database of A-to-I RNA editing. Nucleic Acids Res 42, D109–113.

Ramaswami, G., Lin, W., Piskol, R., Tan, M.H., Davis, C., and Li, J.B. (2012). Accurate identification of human Alu and non-Alu RNA editing sites. Nat Methods 9, 579–581.

Ramaswami, G., Zhang, R., Piskol, R., Keegan, L.P., Deng, P., O’Connell, M.A., and Li, J.B. (2013). Identifying RNA editing sites using RNA sequencing data alone. Nat Methods 10, 128–132.

Rice, G., Patrick, T., Parmar, R., Taylor, C.F., Aeby, A., Aicardi, J., Artuch, R., Montalto, S.A., Bacino, C.A., Barroso, B., et al. (2007). Clinical and molecular phenotype of Aicardi-Goutieres syndrome. Am J Hum Genet 81, 713–725.

Rice, G.I., Kasher, P.R., Forte, G.M., Mannion, N.M., Greenwood, S.M., Szynkiewicz, M., Dickerson, J.E., Bhaskar, S.S., Zampini, M., Briggs, T.A., et al. (2012). Mutations in ADAR1 cause Aicardi-Goutieres syndrome associated with a type I interferon signature. Nat Genet 44, 1243–1248.

Sase, S., Takanohashi, A., Vanderver, A., and Almad, A. (2018). Astrocytes, an active player in Aicardi-Goutieres syndrome. Brain Pathol 28, 399–407.

Schade, M., Turner, C.J., Lowenhaupt, K., Rich, A., and Herbert, A. (1999). Structurefunction analysis of the Z-DNA-binding domain Zalpha of dsRNA adenosine deaminase type I reveals similarity to the (alpha + beta) family of helix-turn-helix proteins. EMBO J 18, 470–479.

Slotkin, W., and Nishikura, K. (2013). Adenosine-to-inosine RNA editing and human disease. Genome Med 5, 105.

Steinman, R.A., and Wang, Q. (2011). ADAR1 isoform involvement in embryonic lethality. Proc Natl Acad Sci U S A 108, E199; author reply E200.

Tan, M.H., Li, Q., Shanmugam, R., Piskol, R., Kohler, J., Young, A.N., Liu, K.I., Zhang, R., Ramaswami, G., Ariyoshi, K., et al. (2017). Dynamic landscape and regulation of RNA editing in mammals. Nature 550, 249–254.

Valente, L., and Nishikura, K. (2007). RNA binding-independent dimerization of adenosine deaminases acting on RNA and dominant negative effects of nonfunctional subunits on dimer functions. J Biol Chem 282, 16054–16061.

van Heteren, J.T., Rozenberg, F., Aronica, E., Troost, D., Lebon, P., and Kuijpers, T.W. (2008). Astrocytes produce interferon-alpha and CXCL10, but not IL-6 or CXCL8, in Aicardi-Goutieres syndrome. Glia 56, 568–578.

Vongpipatana, T., Nakahama, T., Shibuya, T., Kato, Y., and Kawahara, Y. (2020). ADAR1 Regulates Early T Cell Development via MDA5-Dependent and -Independent Pathways. J Immunol 204, 2156–2168.

Wang, K., Li, M., and Hakonarson, H. (2010). ANNOVAR: functional annotation of genetic variants from high-throughput sequencing data. Nucleic Acids Res 38, e164.

Wang, Q., Khillan, J., Gadue, P., and Nishikura, K. (2000). Requirement of the RNA editing deaminase ADAR1 gene for embryonic erythropoiesis. Science 290, 1765–1768.

Wang, Q., Miyakoda, M., Yang, W., Khillan, J., Stachura, D.L., Weiss, M.J., and Nishikura, K. (2004). Stress-induced apoptosis associated with null mutation of ADAR1 RNA editing deaminase gene. J Biol Chem 279, 4952–4961.

Ward, S.V., George, C.X., Welch, M.J., Liou, L.Y., Hahm, B., Lewicki, H., de la Torre, J.C., Samuel, C.E., and Oldstone, M.B. (2011). RNA editing enzyme adenosine deaminase is a restriction factor for controlling measles virus replication that also is required for embryogenesis. Proc Natl Acad Sci U S A 108, 331–336.

Zipeto, M.A., Jiang, Q., Melese, E., and Jamieson, C.H. (2015). RNA rewriting, recoding, and rewiring in human disease. Trends Mol Med 21, 549–559.

